# Melittin-induced membrane disruption in hepatocellular carcinoma involves mtROS/GSDME-mediated pyroptosis

**DOI:** 10.64898/2026.09.05.749568

**Authors:** He Zang, Meng Lu, Nian Fan, Yunzhen Yang, Jianfeng Qiu, Dafu Chen, Rui Guo

**Affiliations:** College of Bee Science, Fujian Agriculture and Forestry University, Fuzhou 350002, China; Department of Pharmacy, The First Affiliated Hospital of Bengbu Medical University, Bengbu 233000, China; National & Local United Engineering Laboratory of Natural Biotoxin, Fuzhou 350002, China; Apitherapy Research Institute of Fujian Agriculture and Forestry University, Fuzhou 350002, China

**Author notes:** These authors contributed equally to this work (He Zang, Meng Lu, Nian Fan, Yunzhen Yang).

**Keywords:** Melittin, Hepatocellular carcinoma, Pyroptosis, Mitochondrial ROS, GSDME

## Abstract

Although melittin is classically regarded as a membrane-lytic peptide, it remains unclear whether its membrane-disruptive effects in tumour cells result exclusively from direct peptide–lipid interactions or are partly generated by pyroptosis. Here, we show that melittin markedly suppresses the viability and clonogenic growth of human Huh7 and murine Hepa1-6 HCC cells and rapidly induces characteristic features of pyroptosis, including cell swelling, balloon-like membrane protrusions, plasma-membrane permeabilization and lactate dehydrogenase release. In Hepa1-6 cells, melittin induced prominent GSDME cleavage and accumulation of its pore-forming N-terminal fragment, whereas GSDMD activation was not detected. Quantitative proteomics and ultrastructural analyses further revealed pronounced mitochondrial perturbation after melittin treatment, accompanied by loss of mitochondrial membrane potential and increased mitochondrial reactive oxygen species (mtROS). Importantly, scavenging mitochondrial oxidants with Mito-TEMPO markedly reduced mtROS accumulation, attenuated pyroptotic membrane damage and partially restored cell viability. Collectively, our findings provide, to our knowledge, the first evidence that melittin induces pyroptotic cell death in HCC cells through a mechanism involving mitochondrial oxidative stress and GSDME processing. These results extend the conventional view of melittin as a direct membrane-lytic peptide and uncover an mtROS/GSDME-associated pyroptotic mechanism underlying its antitumour activity in HCC.

## Introduction

Liver cancer is the third leading cause of cancer-associated mortality worldwide. Hepatocellular carcinoma (HCC) accounts for approximately 90% of primary liver malignancies^(1, 2)^. Treatment options include surgical resection, liver transplantation, radiofrequency ablation, transarterial chemoembolization, targeted therapy and immune checkpoint blockade^(3)^. Even with these treatments, the 5-year overall survival rate remains below 20%^(4)^. Additional therapeutic approaches are therefore needed.

Natural products are an established source of anticancer compounds. Melittin, the main bioactive peptide in bee venom, has antitumour activity in breast cancer^(5)^, non-small cell lung cancer^(6)^, colorectal cancer^(7)^ and ovarian cancer^(8)^. Its amphipathic alpha-helix can insert into lipid bilayers and form transmembrane pores, increasing membrane permeability and causing cell lysis. Melittin also affects intracellular signalling. In breast cancer cells, it inhibits EGFR and HER2 phosphorylation and downstream AKT signalling^(5)^. In ovarian cancer cells, it restricts SREBP1 nuclear translocation, reduces fatty acid synthesis and induces apoptosis^(8)^. The cellular mechanisms engaged by melittin in HCC remain poorly defined.

Pyroptosis is a gasdermin-mediated form of programmed cell death^(9)^. Upstream caspases cleave gasdermins to release pore-forming N-terminal fragments. These fragments oligomerize in the plasma membrane, increase permeability and produce cell swelling and rupture^(10)^. Membrane rupture releases damage-associated molecular patterns (DAMPs) and proinflammatory cytokines^(11, 12)^. Pyroptosis differs from apoptosis, which generally preserves plasma membrane integrity. Pyroptotic cell death can also promote antitumour immune responses^(13)^. Pyroptosis can be initiated by endoplasmic reticulum stress^(14)^, DNA damage^(15)^, lipid metabolic disturbance^(16)^ and mitochondrial reactive oxygen species^(17)^. Several anticancer agents have been reported to kill tumour cells through gasdermin activation^(18, 19)^, suggesting that pharmacological induction of pyroptosis may have therapeutic value.

Whether the membrane-damaging activity of melittin in hepatocellular carcinoma cells arises solely from its direct interaction with lipid bilayers or may also involve the induction of pyroptosis remains unclear. In this study, we therefore investigated the relationship between melittin-induced membrane injury and pyroptotic cell death in HCC cells, with particular attention to gasdermin activation and the associated changes in plasma membrane integrity. By integrating morphological, biochemical and molecular analyses, we sought to determine whether pyroptosis represents a regulated cellular component of melittin-induced membrane permeabilization. This study provides a basis for re-evaluating the antitumour action of melittin beyond its conventional role as a membrane-lytic peptide.

## Materials and methods

### Cell culture and reagents

The human hepatocellular carcinoma cell line Huh7 and the murine hepatocellular carcinoma cell line Hepa1-6 were obtained from the National Collection of Authenticated Cell Cultures (Shanghai, China). Cell-line identity was documented by authentication certificates supplied by the repository. Cells were cultured in Dulbecco’s modified Eagle medium (DMEM) supplemented with 10% fetal bovine serum and penicillin-streptomycin (all from Gibco) at 37 °C in a humidified atmosphere containing 5% COC. Cells were used before passage 30.

Melittin (Selleck Chemicals, cat. no. S9794; purity, 99.79%) was dissolved in sterile water. Dose-response experiments used melittin at 0, 1, 2, 4 or 8 μg/mL. For the single-dose mechanistic experiments, Huh7 and Hepa1-6 cells were exposed to 4 and 2 μg/mL melittin, respectively. Mito-TEMPO was obtained from Selleck Chemicals and used according to the manufacturer’s instructions. In rescue experiments, Mito-TEMPO was maintained in the culture medium throughout the 30-min melittin exposure.

### Cell viability assay

Huh7 or Hepa1-6 cells were seeded in 96-well plates at 2 × 10□ cells per well and allowed to adhere overnight. Cells were exposed to 0, 1, 2, 4 or 8 μg/mL melittin for 30 min, after which the treatment medium was replaced with fresh DMEM. Cell Counting Kit-8 reagent (10 μL per well; Yeasen Biotechnology) was added, and the cells were incubated at 37 °C for 2 h. Absorbance at 450 nm was measured using a microplate reader (Thermo Fisher Scientific). After subtraction of the absorbance from cell-free medium, viability was normalized to that of the corresponding untreated control. For rescue experiments, cells were assigned to control, melittin-alone, Mito-TEMPO-alone or melittin-plus-Mito-TEMPO groups and assayed using the same procedure. Three technical wells were averaged to obtain one value for each biological replicate.

### Colony formation assay

Huh7 or Hepa1-6 cells were seeded in six-well plates at 2 × 10^3^ cells per well in 1 mL of culture medium. After overnight attachment, cells were exposed once to 0, 1, 2, 4 or 8 μg/mL melittin for 30 min. The treatment medium was then replaced with fresh complete medium, and cells were cultured for 7 d without further melittin exposure; the medium was renewed every 24 h. Colonies were fixed with methanol for 10 min, stained with crystal violet (Yeasen Biotechnology) according to the manufacturer’s instructions and imaged. Colony numbers were determined using ImageJ.

### Transwell migration and invasion assays

Migration and invasion were assessed using 24-well cell-culture inserts with 8-μm pores (Biofil, cat. no. TCS003024). For migration assays, 2 × 10□ Huh7 cells or 2 × 10□ Hepa1-6 cells in 200 μL of serum-free medium were added to the upper chamber, and DMEM containing 10% fetal bovine serum was added to the lower chamber. Cells were exposed to 0, 1, 2, 4 or 8 μg/mL melittin during the 24-h migration period. Cells that reached the lower surface of the membrane were fixed with methanol for 10 min, stained with crystal violet and imaged by light microscopy. Migrated cells were counted using ImageJ.

For invasion assays, the upper chamber was coated with Matrigel (Corning) diluted 1:10 in DMEM and incubated at 37 °C for 12 h. Huh7 and Hepa1-6 cells were seeded at 2 × 10□ and 2 × 10□ cells per insert, respectively, and allowed to invade for 24 h in the presence of the indicated melittin concentrations. Fixation, staining, imaging and quantification were performed as described for the migration assay.

### Phase-contrast microscopy

Huh7 and Hepa1-6 cells were exposed to 0, 1, 2, 4 or 8 μg/mL melittin for 30 min and examined immediately by phase-contrast microscopy. Cell swelling and balloon-like membrane protrusions were evaluated in representative fields. For the Mito-TEMPO experiments, cells were exposed to the four treatment conditions described above for 30 min before imaging. Three fields were examined for each biological replicate.

### Scanning electron microscopy

Huh7 and Hepa1-6 cells were left untreated or exposed to 4 and 2 μg/mL melittin, respectively, for 30 min. Samples were fixed and processed using standard procedures for scanning electron microscopy, including dehydration, drying and conductive coating. Cell-surface morphology and pore-like membrane lesions were examined at overview and higher magnifications. Images are representative of three independent biological experiments.

### Lactate dehydrogenase release assay

Huh7 or Hepa1-6 cells were seeded in 96-well plates at 2 × 10□ cells per well and exposed to 0, 1, 2, 4 or 8 μg/mL melittin for 30 min. Culture supernatants were collected, and lactate dehydrogenase (LDH) release was measured using a Cytotoxicity LDH Assay Kit (Dojindo, Japan) according to the manufacturer’s instructions. LDH release was calculated using the controls specified by the manufacturer. For rescue experiments, cells were treated with melittin and/or Mito-TEMPO for 30 min before LDH measurement. Three technical wells were averaged for each biological replicate.

### Propidium iodide uptake assay

Plasma-membrane permeabilization was assessed by propidium iodide (PI) uptake. Huh7 and Hepa1-6 cells were exposed to 0, 1, 2, 4 or 8 μg/mL melittin for 30 min and stained with PI and Hoechst 33342 (Yeasen Biotechnology) according to the manufacturer’s instructions. Fluorescence images were acquired from three fields per biological replicate. The percentage of PI-positive cells was calculated by dividing the number of PI-positive cells by the total number of Hoechst 33342-positive nuclei and multiplying by 100. The same staining and quantification procedure was used after 30 min of treatment with melittin and/or Mito-TEMPO.

### Western blotting

Hepa1-6 cells were exposed to 0, 1, 2, 4 or 8 μg/mL melittin for 30 min, and whole-cell protein extracts were analysed by immunoblotting. Rabbit primary antibodies against GSDMD (Abcam), GSDME (ABclonal) and β-tubulin (Yeasen Biotechnology), together with the corresponding secondary antibody from ABclonal, were used. β-Tubulin served as the loading control. Full-length and N-terminal forms of GSDMD and GSDME were assessed according to their apparent molecular masses. Immunoblots are representative of three independent biological experiments.

### DIA quantitative proteomics and enrichment analysis

Huh7 cells were left untreated or exposed to 4 μg/mL melittin for 30 min, with three independently prepared biological samples analysed per condition. Samples were subjected to data-independent acquisition (DIA)-based quantitative proteomic analysis. Protein identification and relative quantification were performed using the data-processing workflow of the proteomics service provider. Proteins with an absolute log2 fold change greater than 0.263 and *P* < 0.05 were classified as differentially expressed. Differentially expressed proteins were subjected to Kyoto Encyclopedia of Genes and Genomes pathway and Gene Ontology term enrichment analyses. Enrichment results were visualized according to protein number, enrichment P value and, where applicable, rich factor.

### Transmission electron microscopy

Huh7 and Hepa1-6 cells were left untreated or exposed to 4 and 2 μg/mL melittin, respectively, for 30 min. Samples were fixed and processed using standard procedures for ultrastructural analysis, including dehydration, resin embedding, ultrathin sectioning and contrast staining. Sections were examined by transmission electron microscopy. Mitochondrial morphology and cristae integrity were evaluated in overview and higher-magnification images from three independent biological experiments.

### Measurement of mitochondrial membrane potential

Mitochondrial membrane potential was assessed using tetramethylrhodamine methyl ester (TMRM; Selleck Chemicals). Huh7 and Hepa1-6 cells were left untreated or exposed to 4 and 2 μg/mL melittin, respectively, for 30 min. Cells were stained with TMRM according to the manufacturer’s instructions, counterstained with Hoechst 33342 and imaged by fluorescence microscopy. TMRM mean fluorescence intensity was quantified using ImageJ from three fields per biological replicate.

### Measurement of mitochondrial superoxide and Mito-TEMPO rescue

Mitochondrial superoxide was detected using MitoSOX Red (Thermo Fisher Scientific). Huh7 and Hepa1-6 cells were left untreated or exposed to 4 and 2 μg/mL melittin, respectively, for 30 min. Cells were stained with MitoSOX Red and Hoechst 33342 according to the manufacturers’ instructions and imaged by fluorescence microscopy. MitoSOX Red mean fluorescence intensity was quantified using ImageJ from three fields per biological replicate.

To examine the contribution of mitochondrial reactive oxygen species, cells were assigned to control, melittin-alone, Mito-TEMPO-alone and melittin-plus-Mito-TEMPO groups. Mito-TEMPO was applied according to the manufacturer’s instructions and maintained throughout the 30-min melittin exposure. Huh7 and Hepa1-6 cells received 4 and 2 μg/mL melittin, respectively. MitoSOX Red fluorescence, cell viability, cell morphology, LDH release and PI uptake were then assessed using the procedures described above.

### Image analysis

Colony numbers, migrated and invaded cell numbers, fluorescence intensity and the percentage of PI-positive cells were quantified using ImageJ. For imaging assays, three fields were analysed for each biological replicate, and field-level measurements were averaged to obtain one value per replicate. Representative microscopy and immunoblot images were obtained from three independent biological experiments.

### Statistical analysis

All experiments were performed independently three times on different days. Each biological replicate comprised three technical wells or three imaging fields, as appropriate. Technical measurements were averaged within each independent experiment, and the independent experiment was used as the unit of statistical analysis. Data are presented as mean ± s.d., and n denotes the number of independent biological replicates.

The melittin concentration-response data in Fig. 1A, B, D, E, G and H and Fig. 2C and E were analysed by one-way analysis of variance, with each melittin-treated group compared with the corresponding 0 μg/mL control. Comparisons between untreated and melittin-treated cells in Fig. 3F and Fig. 4B were performed using two-sided unpaired Student’s *t*-tests. Data in Fig. 4C, D, F and H were analysed by one-way analysis of variance followed by Tukey’s multiple-comparisons test. *P* ≤ 0.05 was considered statistically significant. Significance is denoted as follows: ns, *P* > 0.05; \**P*≤ 0.05; \*\**P* ≤ 0.01; \*\*\**P* ≤ 0.001; and \*\*\*\**P* ≤ 0.0001.

**Fig. 1.**
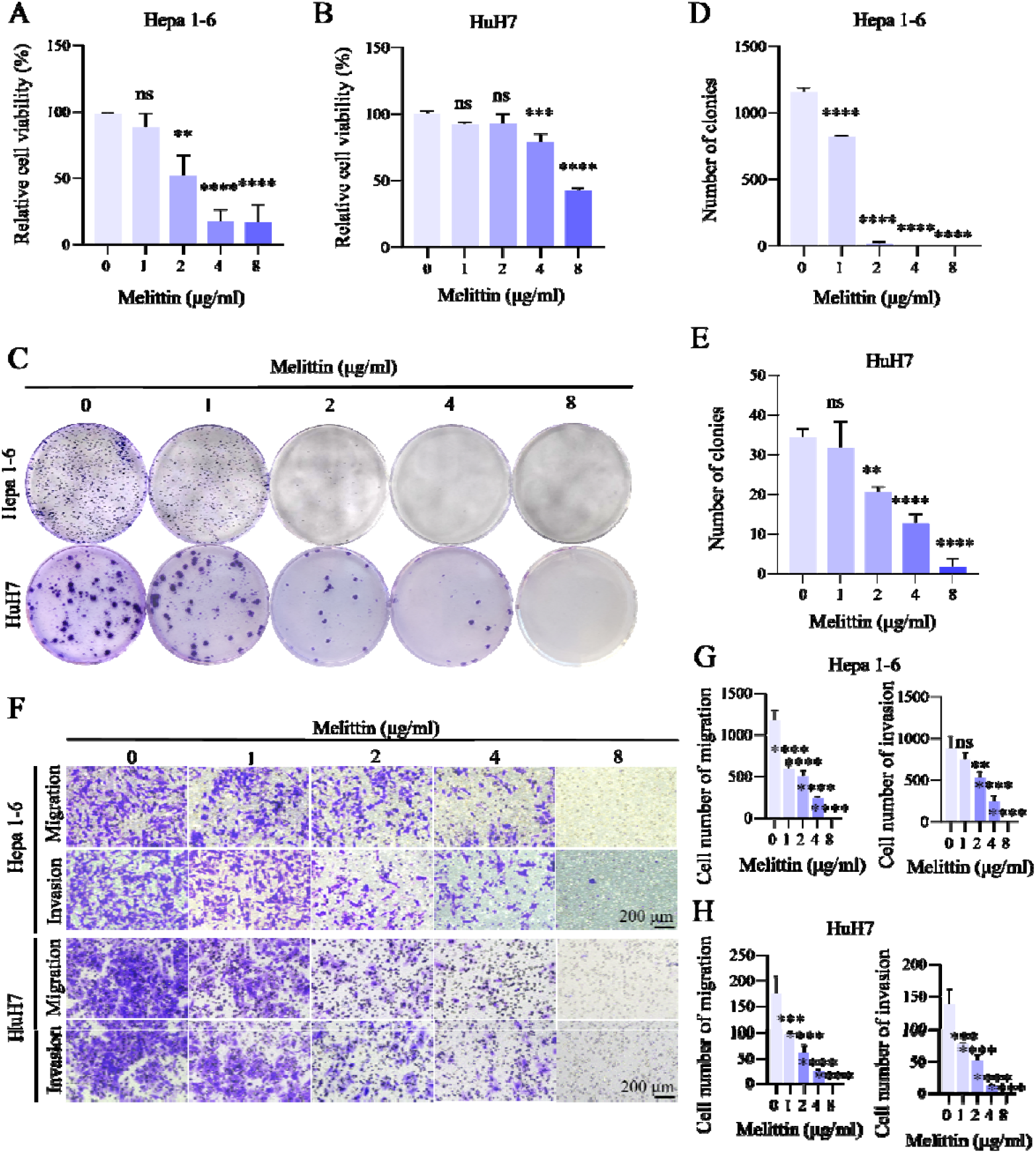
Melittin reduces viability, colony formation and the numbers of migrated and invaded hepatocellular carcinoma cells. **A, B** Relative viability of murine Hepa1-6 (**A**) and human Huh7 (**B**) cells measured using the Cell Counting Kit-8 (CCK-8) assay after 30 min of exposure to melittin at the indicated concentrations, followed by replacement with fresh medium, and normalized to untreated controls. **C** Representative images of crystal violet-stained colonies after a single 30-min exposure to the indicated concentrations of melittin followed by 7 days of drug-free culture. **D, E** Quantification of colony numbers in Hepa1-6 (**D**) and Huh7 (**E**) cells. **F** Representative crystal violet-stained fields from Transwell migration and Matrigel invasion assays performed for 24 h at the indicated melittin concentrations. Scale bars, 200 μm. **G, H** Quantification of migrated and invaded Hepa1-6 (**G**) and Huh7 (**H**) cells. Data in A, B, D, E, G and H are presented as mean ± s.d. from *n* = 3 independent biological replicates; images in C and F are representative of three independent biological experiments. Statistical significance in A, B, D, E, G and H was assessed by one-way analysis of variance (ANOVA), with each melittin-treated group compared with the corresponding 0 μg/mL control. ns, not significant (*P* > 0.05); \**P* ≤ 0.05, \*\**P* ≤ 0.01, \*\*\**P* ≤ 0.001 and \*\*\*\**P* ≤ 0.0001.

**Fig. 2.**
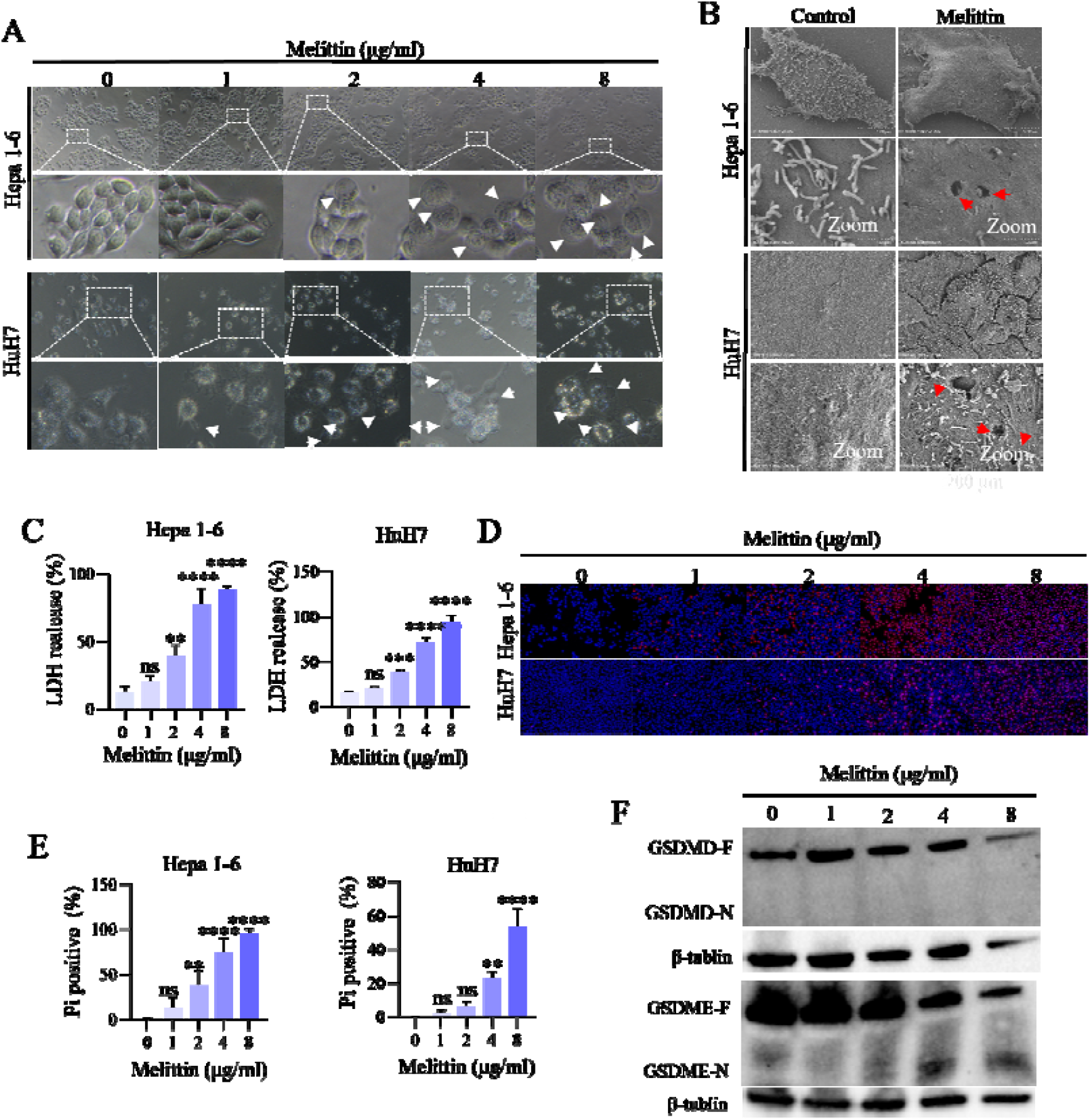
Melittin causes rapid membrane damage with pyroptotic features and GSDME processing in hepatocellular carcinoma cells. **A** Representative phase-contrast images of Hepa1-6 and Huh7 cells exposed to the indicated concentrations of melittin for 30 min. Dashed boxes identify the regions enlarged below, and white arrowheads indicate balloon-like membrane protrusions. **B** Representative scanning electron microscopy images of control and melittin-treated Hepa1-6 and Huh7 cells after 30 min of exposure, with paired higher-magnification views. Red arrowheads indicate pore-like surface lesions. Scale bars are indicated in the images. **C** Lactate dehydrogenase (LDH) release from Hepa1-6 and Huh7 cells after 30 min of exposure to the indicated concentrations of melittin. **D** Representative fluorescence images of propidium iodide (PI; red) uptake in Hepa1-6 and Huh7 cells treated with melittin for 30 min; nuclei are counterstained with Hoechst 33342 (blue). **E** Quantification of PI-positive Hepa1-6 and Huh7 cells. **F** Representative immunoblots of full-length (F) and N-terminal (N) forms of GSDMD and GSDME in Hepa1-6 cells after 30 min of exposure to the indicated concentrations of melittin. Data in C and E are presented as mean ± s.d. from *n* = 3 independent biological replicates; images in A, B, D and F are representative of three independent biological experiments. Statistical significance in C and E was assessed by one-way analysis of variance (ANOVA), with each melittin-treated group compared with the corresponding 0 μg/mL control. ns, not significant (*P* > 0.05); \**P* ≤ 0.05, \*\**P* ≤ 0.01, \*\*\**P* ≤ 0.001 and \*\*\*\**P* ≤ 0.0001.

**Fig. 3.**
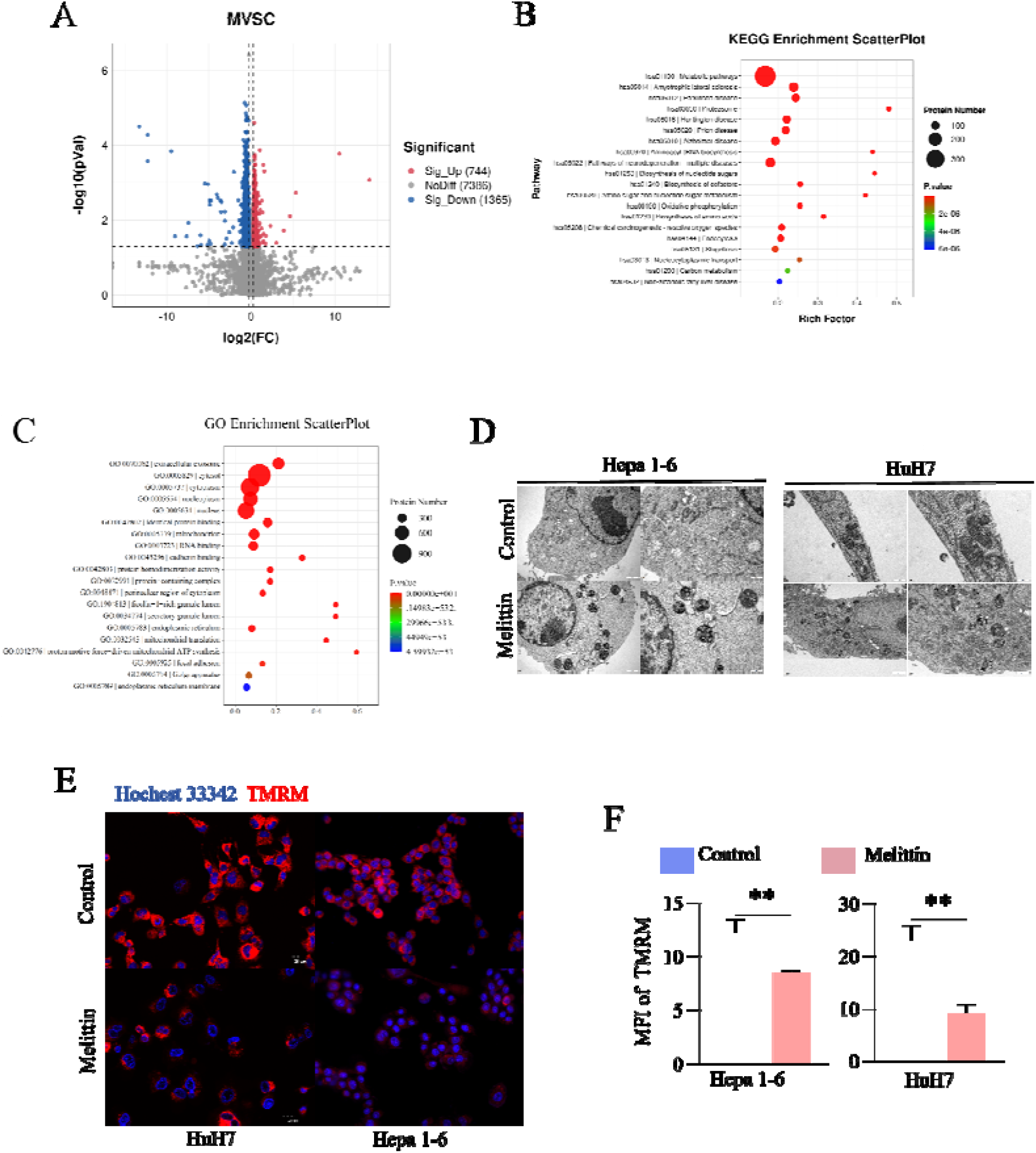
Melittin alters mitochondrial pathways, ultrastructure and membrane potential in hepatocellular carcinoma cells. Control cells received 0 μg/mL melittin; treated Huh7 and Hepa1-6 cells received 4 and 2 μg/mL melittin, respectively. All melittin exposures were performed for 30 min. **A** Volcano plot of quantitative proteomic profiles from control and melittin-treated Huh7 cells. Red, blue and grey points denote 744 upregulated proteins, 1,365 downregulated proteins and 7,386 proteins not meeting the differential-expression threshold, respectively. Dashed lines indicate the stated cut-offs of |log2(fold change)| > 0.263 and *P* < 0.05. **B, C** Kyoto Encyclopedia of Genes and Genomes (KEGG) pathway (**B**) and Gene Ontology (GO) term (**C**) enrichment analyses of differentially expressed proteins. Circle size denotes protein number and colour denotes enrichment *P* value; the x axis in B shows the rich factor. **D** Representative transmission electron microscopy images of Hepa1-6 and Huh7 cells under the indicated conditions, with overview and higher-magnification views. Scale bars are indicated in the images. **E** Representative images of tetramethylrhodamine methyl ester (TMRM; red), a reporter of mitochondrial membrane potential, in Huh7 and Hepa1-6 cells under the indicated conditions; nuclei are counterstained with Hoechst 33342 (blue). Scale bars, 20 μm. **F** Quantification of TMRM mean fluorescence intensity (MFI) in E. All experiments used *n* = 3 independent biological replicates; data in F are presented as mean ± s.d. For each cell line, control and melittin-treated groups in F were compared using a two-sided unpaired Student’s t-test. ns, not significant (*P* > 0.05); \**P* ≤ 0.05, \*\**P* ≤ 0.01, \*\*\**P* ≤ 0.001 and \*\*\*\**P* ≤ 0.0001.

**Fig. 4.**
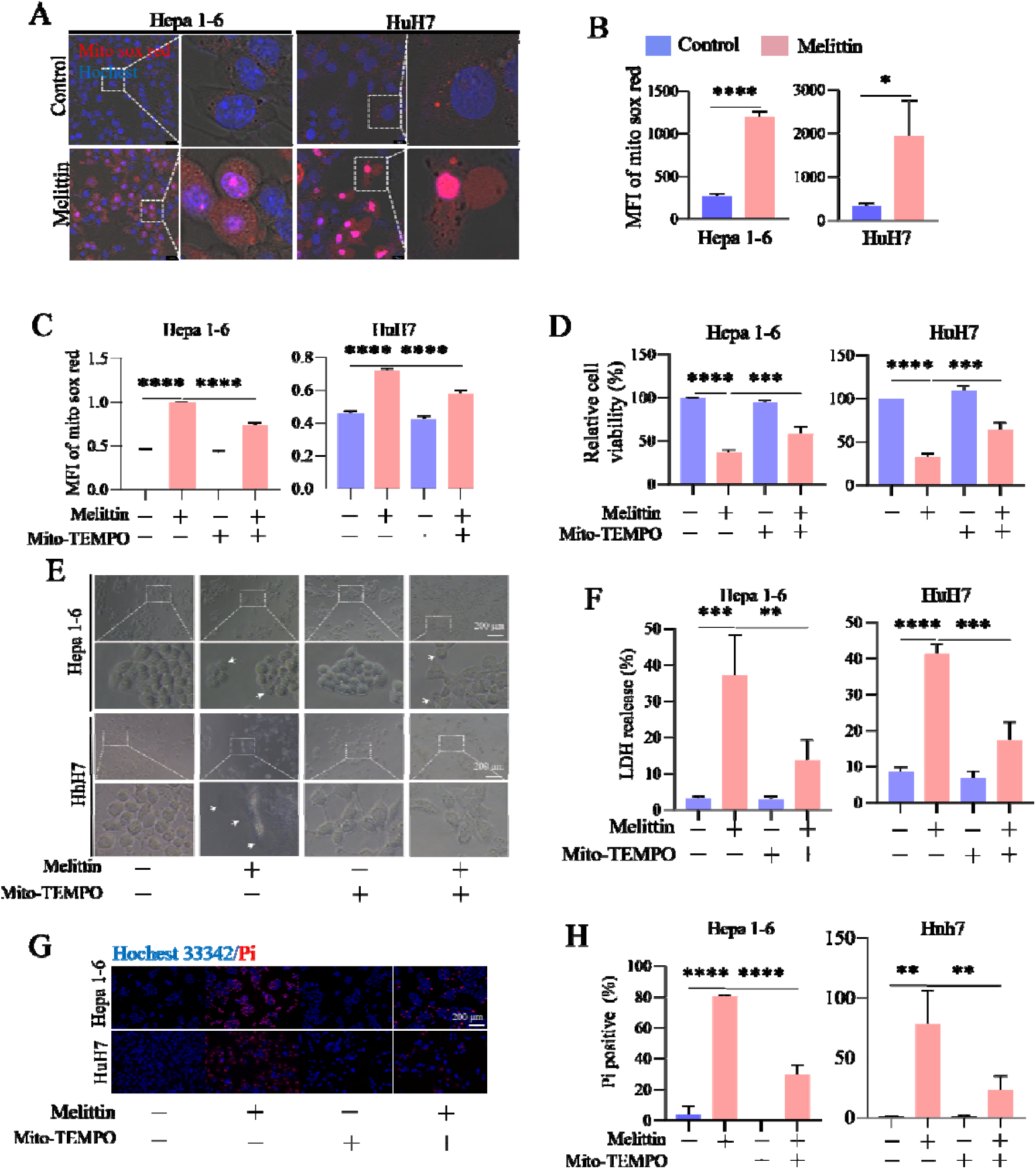
Mito-TEMPO attenuates melittin-induced mitochondrial oxidant signals and membrane damage in hepatocellular carcinoma cells. For experiments involving melittin, control cells received 0 μg/mL; treated Huh7 and Hepa1-6 cells received 4 and 2 μg/mL, respectively. Melittin exposure was 30 min, and Mito-TEMPO was maintained throughout the melittin exposure in the combination-treatment groups. **A** Representative merged transmitted-light and MitoSOX Red fluorescence images of Hepa1-6 and Huh7 cells under control conditions or after melittin treatment. Mitochondrial superoxide is shown in red and nuclei counterstained with Hoechst 33342 are shown in blue; boxed regions are enlarged at right. **B** Quantification of MitoSOX Red mean fluorescence intensity (MFI) in A. **C** MitoSOX Red MFI in cells treated with melittin and/or the mitochondria-targeted superoxide scavenger Mito-TEMPO. **D** Relative cell viability under the same four conditions, measured using the Cell Counting Kit-8 assay. **E** Representative cell-morphology images under the indicated conditions. Boxed regions in the overview images are enlarged below; white arrows indicate cell swelling and balloon-like membrane protrusions. Scale bars, 200 μm. **F** LDH release under the indicated conditions. **G** Representative fluorescence images of PI-positive cells (red); nuclei are counterstained with Hoechst 33342 (blue). Scale bars, 200 μm. **H** Percentage of PI-positive cells in G. Minus and plus signs indicate the absence and presence of the indicated treatment, respectively. Data in B-D, F and H are presented as mean ± s.d. from *n* = 3 independent biological replicates; images in A, E and G are representative of three independent biological experiments. For each cell line, control and melittin-treated groups in B were compared using a two-sided unpaired Student’s *t*-test. Data in C, D, F and H were analysed by one-way ANOVA followed by Tukey’s multiple-comparisons test. ns, not significant (*P* > 0.05); \**P* ≤ 0.05, \*\**P* ≤ 0.01, \*\*\**P* ≤ 0.001 and \*\*\*\**P* ≤ 0.0001.

## Results

### Melittin reduces HCC cell viability and clonogenic growth

We first measured the effects of 0, 1, 2, 4 and 8 μg/mL melittin on Huh7 and Hepa1-6 cells. Cell Counting Kit-8 and colony formation assays showed concentration-dependent reductions in cell viability and colony number in both cell lines (Fig. 1A-E). Increasing melittin concentrations also reduced the numbers of cells that migrated through uncoated Transwell inserts or invaded through Matrigel-coated inserts (Fig. 1F-H).

### Melittin causes membrane damage with pyroptotic features and GSDME cleavage

Melittin produced concentration-dependent cell swelling and membrane blebbing in both HCC cell lines, a morphology consistent with pyroptosis (Fig. 2A). Scanning electron microscopy showed membrane disruption and pore-like surface lesions after treatment (Fig. 2B). Lactate dehydrogenase release and the proportion and fluorescence intensity of propidium iodide-positive cells also increased with melittin concentration (Fig. 2C-E). These measurements show rapid loss of plasma membrane integrity.

Because gasdermin cleavage releases pore-forming N-terminal fragments, we examined GSDMD and GSDME processing in Hepa1-6 cells. Melittin caused concentration-dependent GSDME cleavage and accumulation of GSDME-N, whereas GSDMD-N was not detected (Fig. 2F).

### Melittin alters mitochondrial structure and membrane potential in HCC cells

We next compared the proteomes of untreated and melittin-treated Huh7 cells. At |log2FC| > 0.263 and *P* < 0.05, 744 proteins were upregulated and 1365 were downregulated after treatment (Fig. 3A). Kyoto Encyclopedia of Genes and Genomes and Gene Ontology enrichment analyses associated these changes with mitochondrial processes, including proton-motive-force-driven mitochondrial ATP synthesis and oxidative phosphorylation, and with the chemical carcinogenesis-reactive oxygen species pathway (Fig. 3B,C).

Transmission electron microscopy showed mitochondrial injury and fragmented cristae in melittin-treated HCC cells (Fig. 3D). Tetramethylrhodamine methyl ester fluorescence was also lower after treatment, indicating loss of mitochondrial membrane potential (ΔΨm; Fig. 3E,F). The proteomic and imaging data indicate that melittin rapidly alters mitochondrial structure and function.

### Mito-TEMPO attenuates melittin-induced membrane damage

Melittin increased MitoSOX Red fluorescence in both HCC cell lines, consistent with increased mitochondrial oxidant production (Fig. 4A,B). Mito-TEMPO reduced this signal during melittin exposure (Fig. 4C). It also attenuated cell swelling and membrane blebbing, lowered propidium iodide positivity and lactate dehydrogenase release, and partially restored viability in both cell lines (Fig. 4D-E and Fig. 4F-H). The concordant effects on the mitochondrial oxidant signal and membrane injury support a contribution of mitochondrial oxidative stress to the response.

## Discussion

Melittin has traditionally been regarded as a membrane-lytic peptide whose cytotoxicity mainly reflects direct interactions with lipid bilayers. Biophysical studies have proposed barrel-stave, toroidal-pore, carpet-like and transient-pore models to explain how melittin destabilizes membranes and increases permeability ^(20–26)^. These mechanisms account well for its physical effects on lipid bilayers, but whether membrane rupture in intact tumour cells also involves regulated cell-death machinery has remained less clear. In HCC cells, melittin rapidly induced cell swelling, balloon-like membrane protrusions, plasma-membrane permeabilization, LDH release and pore-like surface lesions, all of which are compatible with pyroptotic membrane rupture. In Hepa1-6 cells, melittin also caused concentration-dependent cleavage of GSDME and accumulation of the pore-forming GSDME-N fragment, whereas GSDMD-N was not detected. Because gasdermin cleavage and release of pore-forming N-terminal fragments are central events in pyroptosis ^(9,10,19)^, these findings support the involvement of a GSDME-associated pyroptotic response in melittin-treated HCC cells. GSDME-mediated pyroptosis has also been linked to tumour suppression and antitumour immunity ^(13)^, raising the possibility that the biological effects of melittin extend beyond direct membrane disruption.

Mitochondrial oxidative stress was closely associated with this response. Proteomic analysis showed marked changes in mitochondrial pathways, including oxidative phosphorylation and proton-motive-force-driven ATP synthesis, together with enrichment of ROS-related processes. These changes were accompanied by mitochondrial structural damage, disruption of cristae, loss of TMRM fluorescence and increased mitochondrial ROS after melittin treatment. Mitochondrial ROS has been implicated in gasdermin-mediated pyroptosis in tumour cells ^(17)^. In our experiments, Mito-TEMPO reduced the MitoSOX signal and, at the same time, alleviated cell swelling, PI uptake and LDH release while partially restoring cell viability. The parallel reduction in mitochondrial ROS and pyroptotic membrane injury suggests that mitochondrial oxidative stress is functionally involved in melittin-induced cell rupture rather than simply arising as a consequence of terminal cell damage. Previous reports that melittin can enter cancer cells through receptor-associated endocytosis ^(27,28)^ are also consistent with the ability of this membrane-active peptide to trigger intracellular responses. Taken together, the mitochondrial damage, mtROS accumulation and GSDME processing observed here support an mtROS/GSDME-linked mechanism underlying melittin-induced pyroptotic membrane rupture.

These findings broaden the current view of melittin cytotoxicity in HCC. Direct disruption of lipid bilayers remains an inherent property of melittin, but our data indicate that membrane permeabilization in intact tumour cells also involves activation of pyroptotic cell death. Mitochondrial dysfunction and mtROS accumulation appear to contribute to this process, together with GSDME processing and pore formation. To our knowledge, this is the first evidence that melittin induces pyroptotic death in cells through an mtROS/GSDME pathway. This mechanism links the classical membrane-lytic activity of melittin with regulated tumour-cell death and provides a new basis for understanding its antitumour effects in hepatocellular carcinoma.

## Data Availability Statement

Data are available from the corresponding author upon request.

## Notes

The authors declare no competing financial interest or other competing interest.

